# Identification of Cell States Using Super-Enhancer RNA

**DOI:** 10.1101/209387

**Authors:** Yueh-Hua Tu, Hsueh-Fen Juan, Hsuan-Cheng Huang

**Author notes:** Correspondence and requests for materials should be addressed to H.F.J. or H.C.H.

## Abstract

A new class of regulatory elements called super-enhancers, comprised of multiple neighboring enhancers, have recently been reported to be the key transcriptional drivers of cellular, developmental, and disease states. Here, we propose to define super-enhancer RNA as highly expressed enhancer RNAs that are transcribed from a cluster of localized genomic regions. Using the cap analysis of gene expression sequencing data from FANTOM5, we systematically explored the enhancer and messenger RNA landscapes in hundreds of different cell types in response to various environments. Applying non-negative matrix factorization (NMF) to super-enhancer RNA profiles, we found that different cell types were well classified. In addition, through the NMF of individual time-course profiles from a single cell-type, super-enhancer RNAs were clustered into several states with progressive patterns. We further investigated the enriched biological functions of the proximal genes involved in each pattern, and found that they were associated with the corresponding developmental process. The proposed super-enhancer RNAs can act as a good alternative, without the complicated measurement of histone modifications, for identifying important regulatory elements of cell type specification and identifying dynamic cell states.

---

Protein-coding genes and other DNA regulatory elements control the amount and activity of proteins in organisms, and constitute the cellular regulatory network. Over the past few decades, transcriptome data has aided in the discovery of numerous facts about gene regulatory networks. However, a systematic understanding of cell differentiation, the development of cancer, and even the dynamic responses of cells to environmental changes remain to be established. Both genetics and epigenetics play important roles in gene regulation. The epigenome may help us extract further knowledge about the interactions with the environment and dynamics of the gene regulatory network.

Enhancers are one of the key links between genetics and epigenetics. Enhancers are activated when transcription factors (TF) bind to them. Subsequently, chromatin modifications direct enhancers to the promoters, and eventually genes are expressed through the actions of TFs. Previously, active enhancers were thought to be tissue-specific and to regulate tissue-specific genes in a spatiotemporal manner^1^. Active enhancer regions are typically decorated by characteristic histone modifications, such as high histone H3 lysine 4 monomethylation (H3K4me1), low histone H3 lysine 4 trimethylation (H3K4me3), and high histone H3 lysine 27 acetylation (H3K27ac). We can identify enhancer loci by detecting histone modifications using chromatin immunoprecipitation sequencing (ChIP-seq); however, prior knowledge is required to design adequate ChIP-seq experiments.

Super-enhancers are large clusters of active enhancers that are densely occupied by TFs, especially master regulators. Super-enhancers are found near key genes in embryonic stem cells; they tend to be cell-specific and regulate the genes essential for cell identity ^2,3^. Super-enhancers differ from typical enhancers in size, TF binding density, and sensitivity to perturbation. Super-enhancers play a role in identifying the key genes for different cell types and are typically identified by the sum of the ChIP-seq signal level of mediators.

In recent years, active enhancers have been found to generate transcripts, called enhancer RNAs (eRNA), which not only promote elongation but also promote chromatin accessibility ^4,5^. Enhancer RNA (eRNA) is a type of noncoding RNA that is generated from the enhancer locus. In depolarized mouse neurons, unusually high amounts of TFs and RNA polymerase II bind to the enhancer locus and bidirectional enhancer RNAs are generated ^6^. Although the function of eRNA remains unknown, eRNA levels can be detected by cap analysis gene expression sequencing (CAGE-seq). Utilizing CAGE-seq, the FANTOM project ^7–9^ analyzed samples from human and mouse and classified into promoter-level expression and enhancer-level expression.

We proposed to define super-enhancer RNA as eRNAs derived from a super-enhancer locus. The expression level of super-enhancer RNA is determined by the sum of eRNA expression levels at a locus. We speculated that super-enhancer RNAs may have the properties of both eRNA and super-enhancers and that they may be positively correlated with the proximal gene and be cell-specific. Further, we explored the classification power of super-enhancer RNAs and identified cell states molecularly using super-enhancer RNA profiles. With knowledge of cell states, we can identify cell behaviors and systemically construct models of cell differentiation or oncogenesis (Fig. 1).

**Figure 1.**
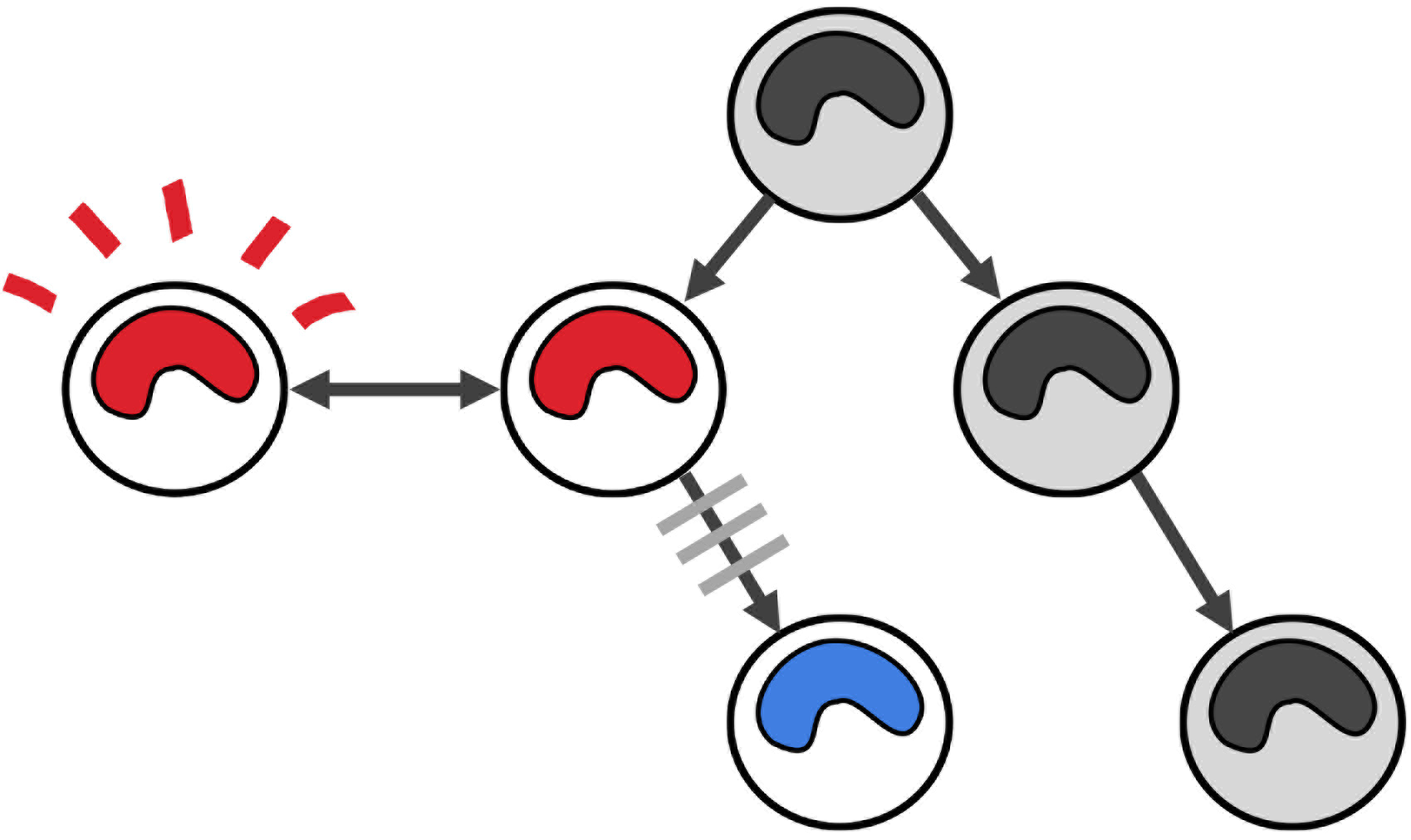
Cell states in the differentiation hierarchy. There should be several cell states during cell differentiation. The stimulated state can also be considered a cell state.

## Results

### Super-enhancer RNA

We obtained enhancer RNA levels from the FANTOM5 project and grouped the enhancer RNAs transcribed from genomic locations within 12.5 kb. We defined the clusters of enhancer RNA transcripts as super-enhancer RNAs and the summed RNA levels as the expression level of the super-enhancer RNA (Fig. 2). Super-enhancer RNA levels and their proximal genes (located within ± 5kb) tended to be positively correlated.

**Figure 2.**
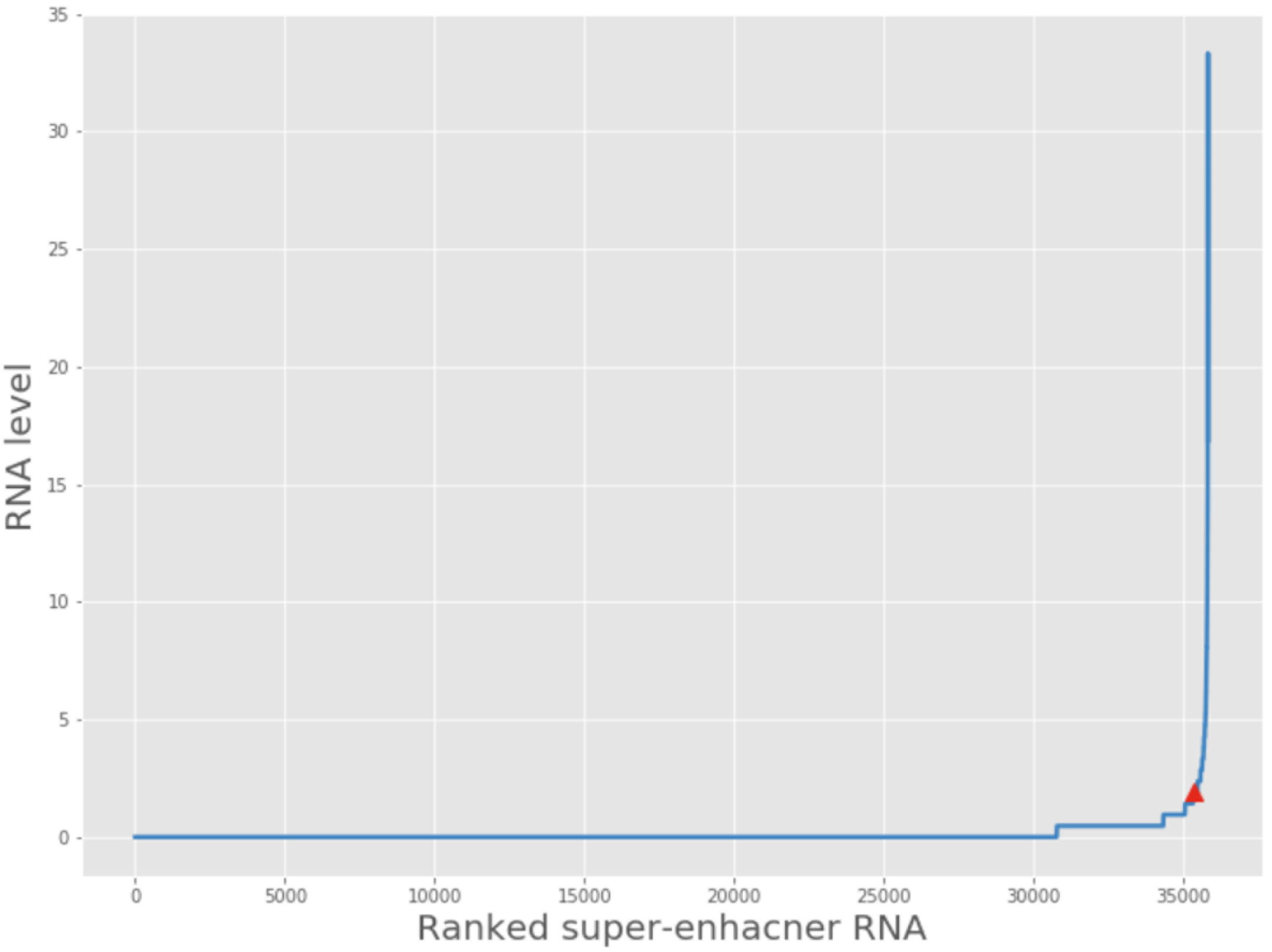
Definition of super-enhancer RNA. The enhancer cluster was merged by summing eRNA levels. Super-enhancer RNAs are defined as the entities to the right of the red triangle.

We compared the super-enhancer loci recorded in dbSUPER database ^10^ with the ones we identified. Regardless of samples, there were 35816 possible unique super-enhancer loci in our data, while dbSUPER presented 65933 possible unique super-enhancer loci. Among them, 50615 (76.8%) unique superenhancer loci from dbSUPER overlapped with ours, while 14351 (40.1%) unique loci from our analyses overlapped with those in dbSUPER. The mismatched loci may arise from the different methods used to measure super-enhancers. Chromatin immunoprecipitation sequencing was performed to identify superenhancers in dbSUPER, while CAGE-seq was performed in the FANTOM5 project.

### Super-enhancer RNAs have higher classification power for cell types than enhancer RNAs

A previous study^11^ has revealed that super-enhancers are cell-specific and we aimed to confirm this using the proposed super-enhancer RNAs. If our superenhancer profiles agree with this cell-specificity, similar samples should be clustered together. First, to establish whether super-enhancer RNAs do have the ability to cluster cells, hierarchical clustering was applied to the time course data (Fig. 3) containing several different cell types under stimulation. Most of the cell types clustered together but there were still some samples mixed in other clusters. Taking a closer look at these specific samples, there were two mixing clusters. One consisted of iPS cells, HES3-GFP embryonic stem cell lines, and H9 human embryonic stem cell lines; the other consisted of mesenchymal stem cells, myoblast to myocyte, aortic smooth muscle cells, and ARPE-19 EMT. All the cell types of the former cluster were stem cell series, while those of the later one were either muscle cell series or the cells belonging to connective tissue.

**Figure 3.**
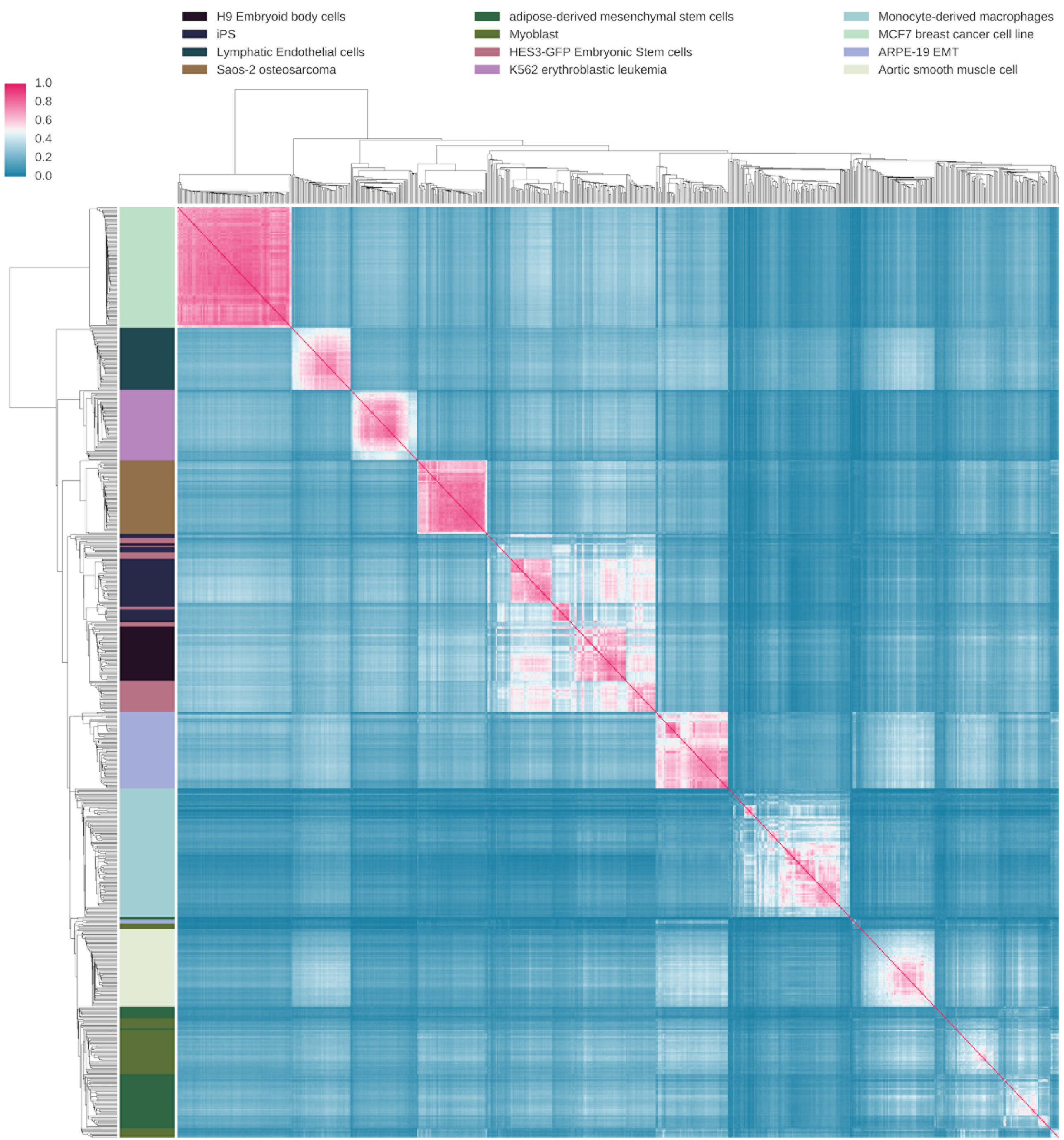
Heat map of the cell type correlation matrix. Pairwise Pearson correlation coefficients were calculated among samples using super-enhancer RNA profiles. Both axes are samples with different cell types. Hierarchical clustering is performed on both axes. Cell types are clustered together, showing the similarities between super-enhancer RNA profiles from different cell types.

To further delve into whether super-enhancer RNAs have classification power for cell types, we performed linear-kernel support vector classification without regularization. We used the same super-enhancer RNA profiles as the features, and the cell types as the prediction target. To compare classification power, we bootstrapped different feature sizes 100 times (Fig. 4). Notably, we found that super-enhancer RNAs had significantly greater classification power for cell types compared with typical enhancer RNAs. The lower the features that were used, the higher the significant difference that was obtained.

**Figure 4.**
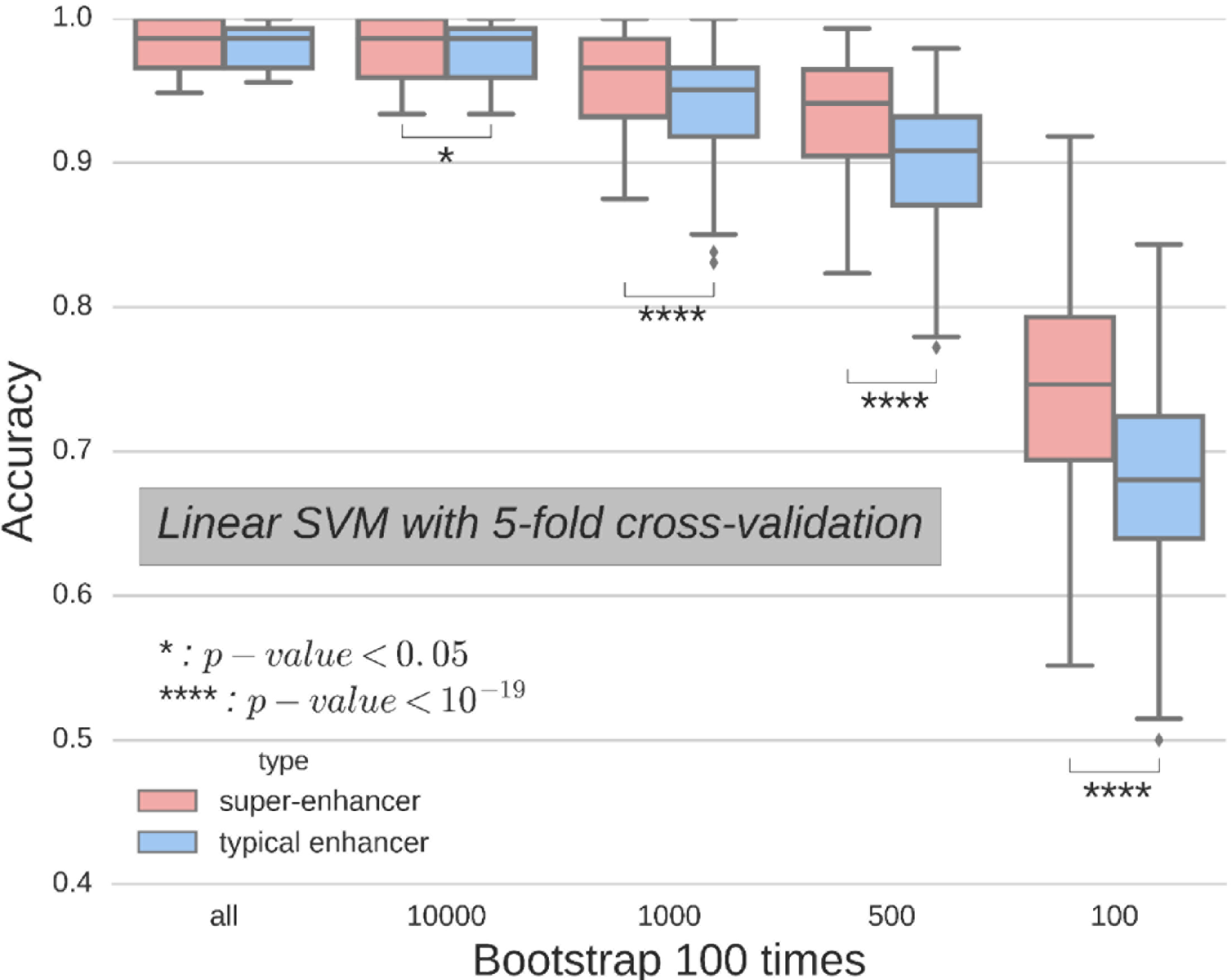
Comparison of classification power between super-enhancer RNA and typical enhancer RNA. All available time course profiles were classified using a linear support vector machine. Different sample sizes were subsampled for each strain and bootstrapped 100 times. Mann-Whitney U tests were performed and p-values are presented.

### Identify cell types using super-enhancer RNA profiles

To identify cell states in the process of differentiation or during dynamic cell responses, we applied non-negative matrix factorization (NMF) on super-enhancer RNA profiles for 12 cell types. If we conceptualize a cell type as a state, NMF may be able to optimally identify different cell types. We found that NMF performs well when *k* was adjusted to close-to but lower than the number of cell types. However, there were still some cell types mixing together, such as iPS cells, HES3-GFP embryonic stem cell lines, and H9 human embryonic stem cell lines, consistent with our earlier results.

### Identifying the cell states of iPS cells differentiating to neural progenitor cells

To identify cell states in the process of cell differentiation, NMF was applied to the time-course experimental dataset of human induced-pluripotent stem cells differentiating to neuronal progenitor cells. In this case, we scanned arbitrary *k* states from 2 to 4 and selected *k* = 3. The super-enhancer profile was factorized into 3 states, and each time point was mapped to one state (Fig. 5). Days 0, 6, and 18 were assigned to the initial state, secondary state, and final state, respectively, while day 12 was selected as the transition state between the secondary and final state.

**Figure 5.**
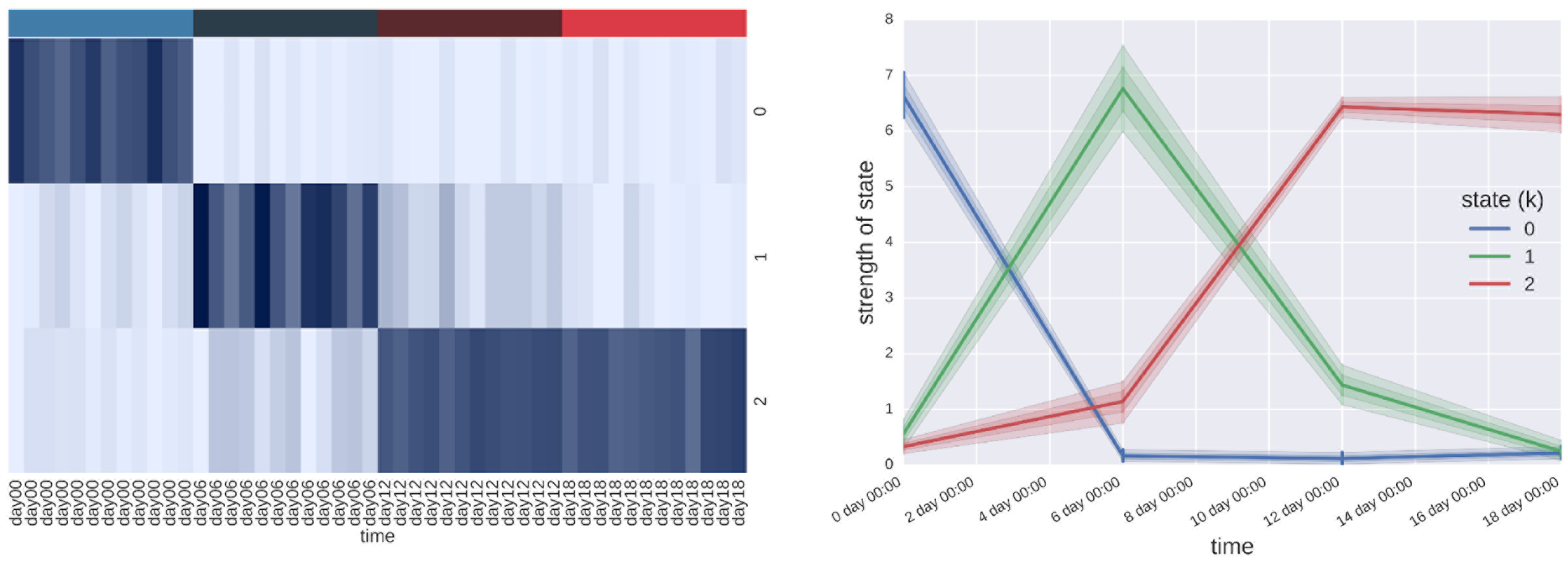
NMF decomposition of time-coursed iPS cell differentiation to neurons. State × sample matrix (left) of the decomposition from the super-enhancer RNA profiles. The x-axis and y-axis reflect the time points and corresponding cell states, respectively, while the darker band represents the greater preference of the cell state. The time series plot (right) was made by collapsing the biological replicates. Progressing patterns, which can be observed in both figures, are interpreted as the transition of cells from one state to another.

To support the identification of cell states, we tested the marker gene expression of stem cells or neurons. We plotted dynamic gene expression over time and found that the expression of neuron progenitor cell markers increased gradually, concurrent with a decrease in the expression of stem cell markers (Fig. 6). The *SOX2* gene is the cell marker gene of both cell types, and we observed that its expression level remained high at each time point.

**Figure 6.**
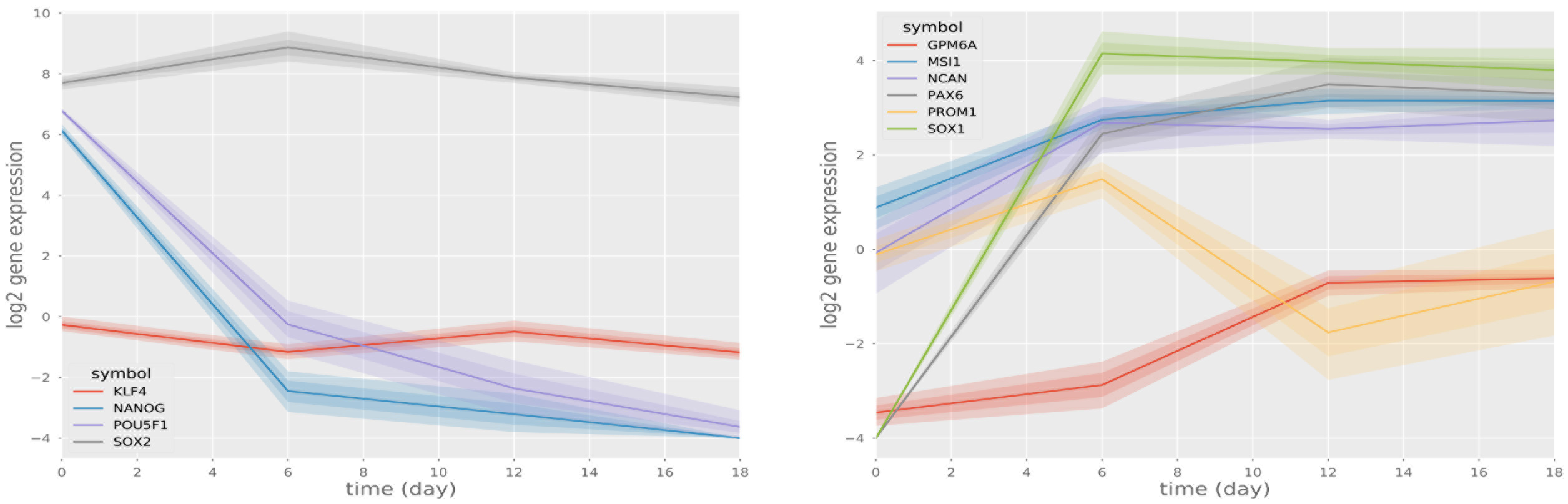
Gene expression of cellular markers. Expression of embryonic stem cell markers (left) declined, while expression of neuron progenitor cell markers (right) rose. The *SOX2* gene acted as the cell marker for both cell types and remained high.

### Functional enrichment analysis of cell states

To further understand the active molecular functions or biological processes in each state, a functional enrichment analysis was applied to SE-assigned genes in each state (Table-1). We chose super enhancers which were enriched in the W matrix (super-enhancer vs. state, *m* × *k*) for each state. SE-assigned genes were mapped and entered into the gene ontology enrichment analysis tool. In the initial state, the RNA biosynthetic process and the metabolic process related GO terms (green) were significantly enriched. The development-related terms (red) were raised to be significantly enriched in the states described below, while the RNA biosynthetic process and metabolic process related terms sank gradually. Terms related to cellular processes (yellow) were randomly distributed in three states, and cellular response-related terms (blue) appeared in the secondary and final states. These findings indicate that cells may actively generate RNAs and maintain their pluripotency initially. Next, cells were treated and induced to differentiate into neurons, while concurrent cellular response processes were activated. The developmental processes elevated gradually and by the end of the experiment the cells remained as neuron progenitors.

**Table 1.** Top 20 enriched functions for SE-assigned genes in each state during iPS cell differentiation to neurons. The differentiation process went from cell state *k*=0 to *k*=2. The *q*-value was adjusted using Benjamini and Hochberg corrections with ConsensusPathDB.

| GO term (k = 0) | q-value | GO term (k = 1) | q-value | GO term (k = 2) | q-value |
| --- | --- | --- | --- | --- | --- |
| regulation of cellular process | 1.3E-12 | regulation of metabolic process | 5.4E-10 | cellular developmental process | 2.0E-07 |
| regulation of RNA metabolic process | 3.8E-11 | regulation of cellular process | 7.3E-10 | regulation of cellular process | 2.0E-07 |
| transcription, DNA-templated | 3.8E-11 | negative regulation of cellular metabolic process | 3.2E-09 | cellular response to chemical stimulus | 5.4E-07 |
| regulation of nucleobase-containing compound metabolic process | 4.4E-11 | regulation of cellular metabolic process | 4.4E-09 | multicellular organismal development | 5.4E-07 |
| RNA biosynthetic process | 1.2E-10 | negative regulation of metabolic process | 7.7E-09 | cell differentiation | 7.4E-07 |
| regulation of gene expression | 1.3E-10 | regulation of macromolecule metabolic process | 7.7E-09 | negative regulation of biological process | 5.1E-06 |
| negative regulation of biological process | 2.4E-10 | regulation of primary metabolic process | 1.5E-08 | system development | 9.4E-06 |
| regulation of metabolic process | 2.4E-10 | negative regulation of cellular process | 1.5E-08 | cellular response to organic substance | 1.3E-05 |
| regulation of macromolecule biosynthetic process | 4.9E-10 | negative regulation of biological process | 1.7E-08 | regulation of signaling | 1.4E-05 |
| cellular response to chemical stimulus | 5.6E-10 | cellular response to chemical stimulus | 1.7E-08 | macromolecular complex subunit organization | 2.1E-05 |
| regulation of nitrogen compound metabolic process | 6.3E-10 | negative regulation of macromolecule metabolic process | 1.7E-08 | muscle structure development | 5.3E-05 |
| regulation of cellular metabolic process | 6.3E-10 | multicellular organismal development | 3.0E-08 | negative regulation of cellular process | 7.2E-05 |
| regulation of primary metabolic process | 8.1E-10 | positive regulation of biological process | 6.0E-08 | positive regulation of biological process | 7.7E-05 |
| cell differentiation | 8.1E-10 | system development | 1.3E-07 | response to organic substance | 7.9E-05 |
| regulation of biosynthetic process | 8.1E-10 | cellular response to organic substance | 2.0E-07 | regulation of metabolic process | 7.9E-05 |
| positive regulation of cellular process | 8.1E-10 | tissue development | 2.4E-07 | regulation of RNA metabolic process | 9.4E-05 |
| negative regulation of metabolic process | 8.1E-10 | regulation of cellular component biogenesis | 4.6E-07 | transcription, DNA-templated | 9.4E-05 |
| regulation of macromolecule metabolic process | 8.2E-10 | regulation of gene expression | 5.0E-07 | regulation of nucleobase-containing compound metabolic process | 9.4E-05 |
| positive regulation of biological process | 8.8E-10 | cellular developmental process | 7.9E-07 | regulation of macromolecule biosynthetic process | 9.4E-05 |
| cellular developmental process | 8.8E-10 | positive regulation of cellular process | 1.0E-06 | regulation of gene expression | 1.0E-04 |

metabolic process
signal transduction, cellular process
development
cellular response

### Cell states in macrophages

We further performed NMF on the macrophage response to the LPS experiment from the FANTOM project (Fig. 7). In the original experiment, macrophages were treated with LPS, which is the material on the surface of gram-negative bacteria, and expression profiles were measured from 0 to 48 hours. LSP should stimulate the innate immune pathways in macrophages and we observed this pattern of activation in the H matrix from the NMF analysis. The cell state (*k*=1) is the active state and the alternative (*k*=0) is the inactive state; we could observe a peak at the early stage.

**Figure 7.**
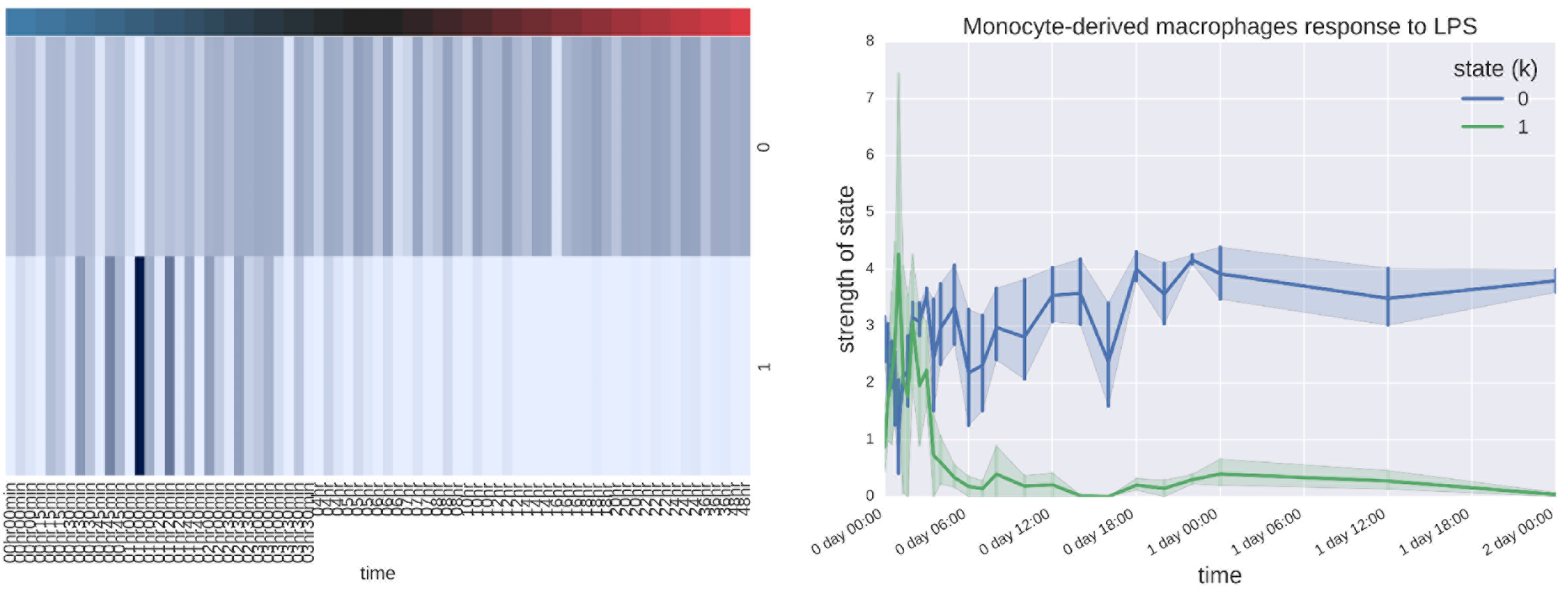
NMF decomposition of the time-coursed macrophages response to the LPS experiment. State × sample matrix (left) of decomposition from super-enhancer RNA profiles. The x-axis and y-axis stand for the time points and cell states, respectively, while the darker band represents the cell state’s greater preference. The time series plot (right) was made by collapsing the biological replicates. The stimulation peak, which can be observed on both figures, was interpreted as cells that are stimulated and transition into an active state.

## Discussion

CAGE-seq, which was used in the FANTOM5 project, targets the 5’ cap of transcripts, which is beneficial for the detection of eRNA. Bi-directional enhancer RNAs do not process the post-transcription modification of RNA splicing and polyadenylation as messenger RNA, but do possess 5' caps. On the other hand, one limitation of RNA-seq is that it is not able to detect eRNAs. Instead, CAGE-seq is necessary to quantize the activity of enhancers and super-enhancers, which enables comparison with gene expression.

We have proposed using super-enhancer RNA to identify cell states. Initially, we found a positive correlation between super-enhancer RNA and the SE-assigned gene. Additionally, cell types were well-classified by super-enhancer RNAs. Super-enhancer RNA may inherit its positive correlation with the expression of the nearest gene and its cell-specificity from eRNA and super-enhancers, respectively. We have further shown the classification power of super-enhancer RNA profiles by training a linear support vector machine. Instead of a sophisticated and powerful classification model, the simple linear classification model demonstrated the linear-separability of super-enhancer RNA profiles. With regards to directions for future investigation, we now aim to evaluate whether super-enhancer RNA profiles could provide further information beyond cell identity.

Previously, researchers have identified cell types from cell morphology and molecular markers. Here, we demonstrated an approach that distinguishes cell types based on molecular configurations using NMF to identify cell types and states. NMF can be conceptualized as the linear combination of nonnegative column vectors. Interpretation of the matrix decomposition depends on determination of the input matrix. For different cell-type profiles, cell types are clustered together into k clusters; for cell differentiation profiles, similar molecular states in progress are clustered together, which demonstrate the pattern of progression from one cell type to another. NMF is an excellent tool for excavating the latent variables in cell profiles. Yet one limitation of the method is that the determination of k must be manually instituted. Optimal results are attained if hypotheses are strong and data quality is high.

Super-enhancers appear proximal to key identity genes in different cell types ^3^. In many cancer cells, super-enhancers emerge near the oncogenic drivers ^12^, thus, the regulatory patterns may be similar in normal cell types and cancer cells. Both oncogenic driver genes and master regulators are the regulators that govern and maintain cell identities. If regulators are down-regulated, cells lose their properties and behaviors. The core regulatory circuitry consists of master regulators which are auto-regulated and, in turn, regulate each other forming a regulatory clique ^13^. Super-enhancers act as guides for cell-specific genes and master regulators. Further, genome-wide association studies (GWAS) show that most disease-associated single nucleotide polymorphisms (SNPs) are located in the noncoding region, especially within enhancers ^8^. One recent study supports the hypothesis that the formation of super-enhancers is not only related to cell identity, but is also related to changes in cell state ^11^. Super-enhancers are the key switches in the gene regulatory network and the link to disease.

Another recent study ^14^ has revealed the relationship between super-enhancers and pioneer factors. Pioneer factors are informally defined as the first transcription factor which promotes untying the wrapped histone that releases the bare DNA. After pioneer factors reach the target histone, super-enhancers are established gradually by a selection of key transcription factors. During this process, both super-enhancers and the gene regulatory relationship are remodeled. The removal of old super-enhancers and establishment of new superenhancers changes the cell identity and the transcription of master regulators ^14^; chromatin modification is updated later ^15^. Identification of cell states may be the key step to identifying the progression of cell and pioneer factors.

The super-enhancer RNAs we proposed here may be a new means of measuring the activity of super-enhancers; they act as a good alternative for the classification of cell type specification, and do not require the complicated measurement of histone modifications by ChIP-seq. Super-enhancer RNA profiles provide the opportunity to identify cell types or states. NMF is a good method for decomposing large biological data to reveal interpretable latent variables. We further plan to investigate the dynamics of cell development, cell response, and cancer development based on these findings.

## Methods

### FANTOM data

We obtained gene expression data and enhancer RNA level data from FANTOM5 and downloaded it using the FANTOM5 Table Extraction Tool (http://fantom.gsc.riken.jp/5/tet/#!/search/hg19.cage_peak_ann.txt.gz). We selected the &“Human Phase 1 and 2&” option in the dataset and downloaded whole read counts and RLE normalized expression data. We downloaded enhancer RNA levels from (http://fantom.gsc.riken.jp/5/datafiles/latest/extra/Enhancers/) and selected the normalized enhancer RNA expression table (human_permissive_enhancers_phase_1_and_2_expression_tpm_matrix.txt.gz).

### Data Preprocessing

#### Mapping enhancers to proximal genes

We parsed all enhancer and gene locus information and established putative regulatory relationship between enhancers and their proximal genes located within ±5kb from their midpoint. If there was no gene located within ± 5kb of an enhancer, we assigned the nearest gene to it.

#### Identifying super-enhancers

We stretched enhancers to make stitched enhancers using the midpoint of each enhancer locus as a reference. Enhancers located within 12.5 kb were combined into stitched enhancers. To avoid the overlapping of gene loci, if there were gene loci located between two combining enhancers, we retained the two enhancer loci as separate. Expression levels of eRNA were calculated to identify supere-enhancers. All stitched enhancer loci were ranked by the sum of their eRNA levels, and super-enhancers were defined as those which had summed expression levels higher than the reflection point of the eRNA distribution curve (Fig. 2).

#### Mapping super-enhancers to the nearest gene

We parsed the super-enhancer loci and gene loci for location information, and assigned each super-enhancers to its nearest gene according to the end-point of the super-enhancer locus.

### Comparison to known super-enhancers

dbSUPER is an integrated and interactive database of super-enhancers, which contains 82234 super-enhancers from 102 human and 25 mouse tissue/cell types. All dbSUPER human super-enhancer loci were obtained from the website (http://bioinfo.au.tsinghua.edu.cn/dbsuper/). Then, all dbSUPER superenhancer loci and our super-enhancer RNA loci were parsed and duplicate loci were removed. Overlapping analyses were applied to both sets of super-enhancer loci. In each comparison, the length of the overlapping region was calculated then divided by the length of each locus. If the overlapping rate was larger than 50%, the two loci were considered to be overlapped.

### Heat map of cell types

All super-enhancer loci were used to perform hierarchical clustering. Time course expression data were used and were transformed to a log scale. To avoid imbalanced training, we ruled out cell types which had a sample size of less than 20. We replaced the negative infinity value in the log-scale with the minimum of the whole expression matrix. We performed a Pearson correlation matrix on the log-expression profile, then performed hierarchical clustering on the correlation matrix using Euclidean distance metric and single method algorithm.

### Cell type classification

All super-enhancer and enhancer expression profiles were used. After ruling out small sample size (< 20) cell types, logarithmic scale transformation, and replacing the negative infinity value with the minimum of the matrix, we performed a classification analysis on cell types. Support vector classification was applied to the log-expression profile with a linear kernel and five-fold cross-validation. Cross-validation scores were collected as accuracies, and randomly sampled 100 times for a different number of loci in each analysis. Mann-Whitney U tests were performed on each analysis.

### Non-negative matrix factorization

We transformed the super-enhancer RNA data matrix into a logarithmic scale and replaced the negative infinity value with the minimum value of the matrix. Shifting minimum to zero to keep values non-negative, we applied NMF without regularization using the scikit-learn Python package. The matrix was factorized into *W* (super-enhancer RNA vs. state, *m* × *k*) and *H* matrices (state vs. sample, *k* × *n*). With regards to cell types, all available time course super-enhancer RNA profiles were used and cell types were later labeled, but sample sizes smaller than 20 omitted to avoid imbalanced model training. To avoid the local optimal solutions, we repeated the same process with 200 random initial conditions and selected the best one evaluated using their silhouette scores. We evaluated the *H* matrix of each NMF model by assigning the highest preference state to each sample. With regards to cell states, the time course of single experiment superenhancer profiles, e.g., iPS cell that differentiated to neurons, were used and time points were later labeled.

### Functional enrichment analysis

Highly enriched super-enhancer RNAs in each state were selected from the factorized *H* matrix (state vs. sample). SE-assigned genes for each state were obtained using the method described above and the human gene function annotation, ConsensusPathDB-human (http://cpdb.molgen.mpg.de/), was used for the functional enrichment analysis. List files of the HGNC gene symbols of each state were uploaded to the website and the top 20 significant gene ontology (GO) terms from levels 3~5 for each state were obtained.

### Cell marker genes

>We obtained seven neuron progenitor cell marker genes (*SOX2*, *PAX6*, *MSI1*, *PROM1*, *NCAN*, *SOX1*, *GPM6A*) (Tian *et al*, 2013) and four stem cell marker genes (*POU5F1*, *SOX2*, *NANOG*, *KLF4*) and plotted the time series of gene expression on a log scale.

## Acknowledgements

This work was supported by Ministry of Science and Technology, Taiwan (MOST 103-2320-B-010-031-MY3, 104-2628-E-010-001-MY3, and 105-2320-B-002-057-MY3) and National Health Research Institutes (NHRI-EX105-10530PI).

## Author contributions

YHT, HFJ and HCH conceived and designed the study. YHT implemented the method and performed data analysis. YHT, HFJ and HCH interpreted the data. YHT, HFJ and HCH wrote the manuscript. HFJ and HCH supervised the study. All authors read and approved the final manuscript.

